# Fast Memory Integration Facilitated by Schema Consistency

**DOI:** 10.1101/253393

**Authors:** Qiong Zhang, Vencislav Popov, Griffin E. Koch, Regina C. Calloway, Marc N. Coutanche

## Abstract

Many of our everyday decisions are based not only on memories of direct experiences, but on memories that are integrated across multiple distinct experiences. Sometimes memory integration between existing memories and newly learnt information occurs rapidly, without requiring inference at the time of a decision. Such fast memory integration is known to be supported by the hippocampus but not the neocortex. In this study, we explore an alternative mechanism of fast memory integration, through related prior knowledge (i.e., schema), which is associated with neocortical learning. Paired associates were selected to be schema consistent or inconsistent, and confirmed with a latent semantic analysis of text corpora. We observed that after enabling fast learning by using material that is consistent with a schema, faster memory integration can occur. This result suggests that the hippocampus-mediated integration of new information is not the only available mechanism that supports fast memory integration.

## INTRODUCTION

Memory is essential in guiding everyday decisions. These decisions are not only based on memories of direct experiences alone, but also rely on knowledge generalized across multiple distinct events. One process that supports such generalization is memory integration. Effective memory integration transforms distinct, but overlapping past experiences into a cohesive representation (Eichenbaum, 2000; Gluck & Myers, 1993), on which one can base novel judgments later (Zeithamova, Dominick, & Preston, 2012). These novel memory decisions can be supported either by direct integration during the encoding of overlapping elements (i.e., “integrative encoding”; Shohamy and Wagner, 2008) or by inferring the relations between elements during retrieval (i.e., “logical inference”; Dusek and Eichenbaum, 1997; Greene, Gross, Elsinger, & Rao, 2006). Research has suggested that both mechanisms depend on hippocampal integration, and that neocortical generalization plays a limited role in the integration process (Shohamy and Wagner, 2008; Greene et al., 2006; Heckers et al. 2004; Preston et al., 2004). In this study, we investigated the possibility that other mechanisms –inspired by learning mechanisms that are neocortically based– can enable fast memory integration. We examined how schema-consistent and schema-inconsistent information (i.e., consistency of new information with prior knowledge) affect inference and integration in a paired-associate inference task.

Memory integration is most commonly examined with the associative inference paradigm. In this task, participants learn separate events with overlapping components (e.g. A-B and B-C), and later have to infer the relations between elements that have not been experienced together but are indirectly associated (e.g., A-C; Shohamy and Wagner, 2008; Myers et al., 2003; Preston, Shrager, Dudukovic, & Gabrieli, 2004). Two different mechanisms have been proposed to explain how participants make such indirect inferences. The first mechanism, integrative encoding, is a fast method of memory integration that takes place during encoding through dynamically shifting between encoding and retrieval states of the hippocampus. This proposed mechanism is supported by experimental studies (Shohamy and Wagner, 2008), and is consistent with computational theories (Hasselmo and McClelland, 1999; Hasselmo, Schnell, & Barkai, 1995). The second mechanism, logical inference, does not involve directly encoding an integrated memory (i.e., A-C), and instead infers the relationship between A and C after retrieving separate memories of A-B and B-C (Dusek and Eichenbaum, 1997; Greene et al., 2006).

There is extensive evidence that both mechanisms are supported by the hippocampal system. During integrative encoding, related prior experiences that overlap with the newly encoded information are reactivated in the hippocampus (Schlichting and Preston, 2015). Memory integration then takes place at the time of learning, supported by the integration of new experiences into existing memory networks by the hippocampus (Shohamy and Wagner, 2008; Zeithamova and Preston, 2010). Evidence for this comes from both non-human animal and human studies. Animal studies have demonstrated that the hippocampus can encode similarities between distinct events (Eichenbaum et al., 1999; Wood, Dudchenko, & Eichenbaum, 1999; Singer et al., 2010), and can reactivate traces of prior events, when learning new information (Karlsson and Frank, 2009). More direct evidence comes from recent neural imaging studies with humans – hippocampal activity during encoding predicts subsequent performance in memory integration (Shohamy and Wagner, 2008; Zeithamova and Preston, 2010; Schlichting, Zeithamova, & Preston, 2014). Similarly, it has been suggested that the hippocampus also supports flexible retrieval of component memories (A-B and B-C) during logical inference (Greene et al., 2006; Heckers et al. 2004; Preston et al., 2004).

Thus, current theories of memory integration posit that both integrative encoding and logical inference are mediated by the hippocampus, and that the neocortex plays a limited role in this rapid integration process. According to the complementary learning systems theory (Marr, Willshaw, & McNaughton, 1991; McClelland, McNaughton, & O’Reilly, 1995), the brain keeps two separate memory stores to avoid interference between new information and existing memories. Initial learning takes place in the temporal store supported by the hippocampus. Through system consolidation involving both time and sleep, newly learnt information gradually transfers to a more permanent store supported by the neocortex (Zola-Morgan and Squire, 1990; Frankland and Bontempi, 2005; Born and Wilhelm, 2012). This standard view suggests that integrating A and C into one representation shortly after learning A-B and B-C cannot depend on neocortical areas.

However, recent findings suggest that system-level consolidation can take place rapidly. Newly learnt information that is consistent with pre-existing knowledge (i.e. schema) becomes independent of the hippocampus (Tse et al., 2007, 2011), with the medial prefrontal cortex (mPFC) shown to inhibit the hippocampal binding (Bein, Reggev & Maril, 2014; van Kesteren, Ruiter, Fernandez, & Henson, 2012; van Kesteren et al., 2013). This effect was also confirmed by recent simulations under the complementary learning systems theory, in which assimilating schema-consistent knowledge occurred rapidly without interference with existing neocortical representations (McClelland, 2013). Recent studies have also shown that word-concept associations can become rapidly integrated into lexical memory if related knowledge is accessed during encoding through a “fast mapping” procedure (Coutanche and Thompson-Schill, 2014; Coutanche and Thompson-Schill, 2015). This rapid integration draws on neocortical systems (Merhav, Karni, & Gilboa, 2015), without requiring the hippocampus (Sharon, Moscovitch, & Gilboa, 2011), and might share mechanisms with the rapid learning that is induced by a schema (Coutanche and Thompson-Schill, 2015).

It is currently unclear whether memory integration in the associative inference task can be supported by neocortical mechanisms. If associate pairs A-B and B-C are linked through a schema, it might be possible for neocortical mechanisms to also contribute to integrative encoding (i.e., associating A and C). In this study, we examine the possibility for rapid memory integration in an associative inference task (e.g., Shohamy et al.; 2008), in a way that is more consistent with neocortically based mechanisms. We will compare memory integration for schema consistent and schema inconsistent pairs. Participants learn associative word pairs in the format of person-location, with some pairs that are schema-consistent (e.g. teacher-classroom, classroom-student) and others that are schema-inconsistent (e.g. baker-theater, theater-hiker). Given the evidence that episodic associations between schema-consistent pairs are learned neocortically, and that hippocampal involvement is suppressed in those cases (Bein et al, 2014; van Kesteren et al., 2012; van Kesteren et al., 2013), if the neocortex <u>cannot</u> support memory integration, relatedness inference decision should be worse for schema-consistent pairs (i.e., teacher-student) than for schema-inconsistent pairs (i.e., baker-hiker). In contrast, if rapid integration in the associative inference task can be supported by the neocortex, and given that semantically related pairs are easier to learn, we should observe easier memory integration for schema-consistent pairs. The extent of memory integration due to integrative encoding from that due to logical inference can be distinguished by introducing two levels of memory integration. In addition to generalizing A-B and B-C to A-C (i.e., 1-link integration), we also test generalizing A-B, B-C and C-D to A-D (i.e., 2-link integration). If participants respond to tests based on integrative encoding, the response time (i.e., RT) should reflect a direct retrieval that is independent of the number of links. If participants respond to tests based on logical inference, the RT should be dependent on the number of links, as the inference process involves cognitively traversing every link.

## METHOD

### Participants

Thirty-five participants (17 females; mean (M) age = 20.7 years, standard deviation (sd) = 3.0; English speakers without a learning or attentional disorder) contributed to the study. Informed consent was obtained for each participant prior to beginning the study. Upon completion, participants were compensated through course credit or payment for their time. The University of Pittsburgh Institutional Review Board approved all procedures. Eight participants were excluded from the analysis – five participants did not reach criterion for at least half of the studied pairs (see Procedure); three participants showed chance performance during forced-choice testing. Exclusion criteria were established prior to the start of data collection.

### Materials

To test the effects of schema consistency on memory integration, we implemented a 2 (schema consistency: consistent vs. inconsistent) x 2 (linked pairs: one-link integration vs. two-link integration) within-subjects design. Schema consistency in the present study is based on the association between a person (e.g., teacher) and a location (e.g., classroom). Within the experiment, one ‘set’ consists of three word-pair associations (i.e., person1-location1, location1-person2, person2-location2). There are ten sets within each schema condition, resulting in 60 unique word-pair associations to be studied. Sets are trained in the study phase of the experiment.

**Table 1.**
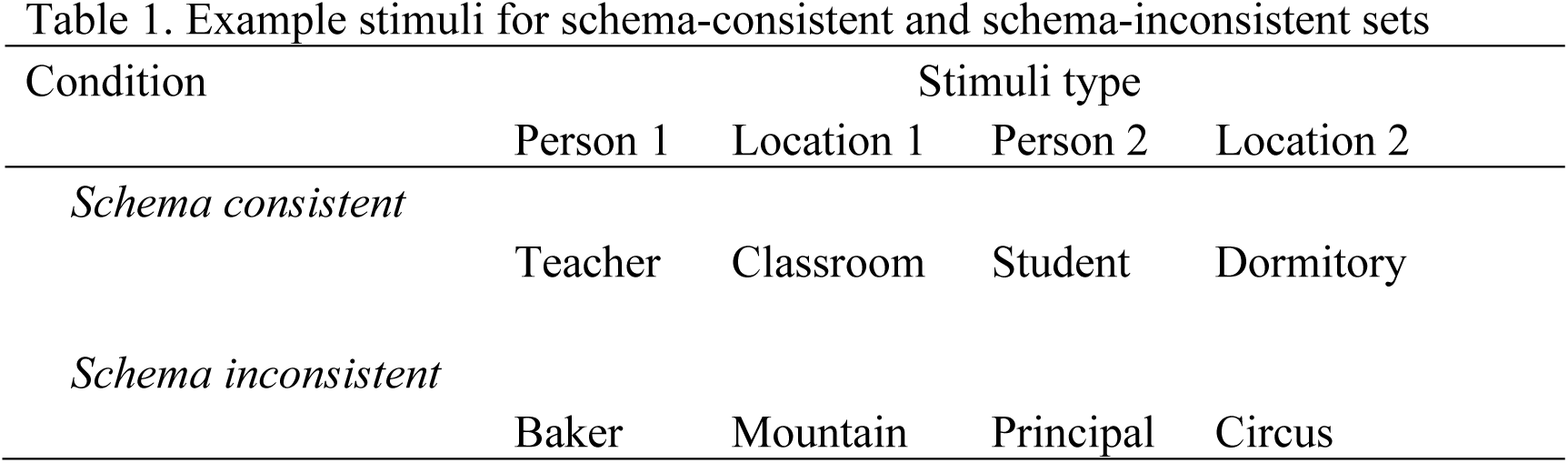
Example stimuli for schema-consistent and schema-inconsistent sets

Latent semantic analysis (LSA; gathered from <u>http://lsa.colorado.edu/</u>) was used to confirm schema consistency. LSA can be used to measure word associations based on their co-occurrence within large corpora (Landauer & Dumais, 1997). LSA values range from −1 to 1, with higher values indicating stronger semantic associations. Word pairs in the schema-consistent condition (*M_LSA_* = .42, *SE* = .04) had higher LSA scores than word pairs in the schema-inconsistent condition (*M_LSA_* = .06, *SE* = .01*; t*(58) = 8.59, *p* < .001).

In the test phase, participants are tested on their memory for the pairs in the studied sets and the linked pairs. One-link integration is the association between person1 and person2. Two-link integration is the association between person1 and location2.

### Procedure

All Experimental procedures were created and presented using PsychoPy2 Experiment Builder, v1. 84. 2 (Peirce 2007; 2009).

### Study phase

#### Learning task

During the study phase of the experiment, participants were presented with two words (a person and a location) and instructed to remember the pairing. To help them remember the words, participants had to decide how likely it would be to see the person/profession in the paired location, on a 4-point scale (very likely, somewhat likely, somewhat unlikely, very unlikely). Each trial began with a fixation cross presented in the middle of the screen for 0.5 s., followed by the word pair, which remained on the screen for 3.5 s. regardless of when participants responded. The procedure is shown in Figure 1.

**Figure 1.**
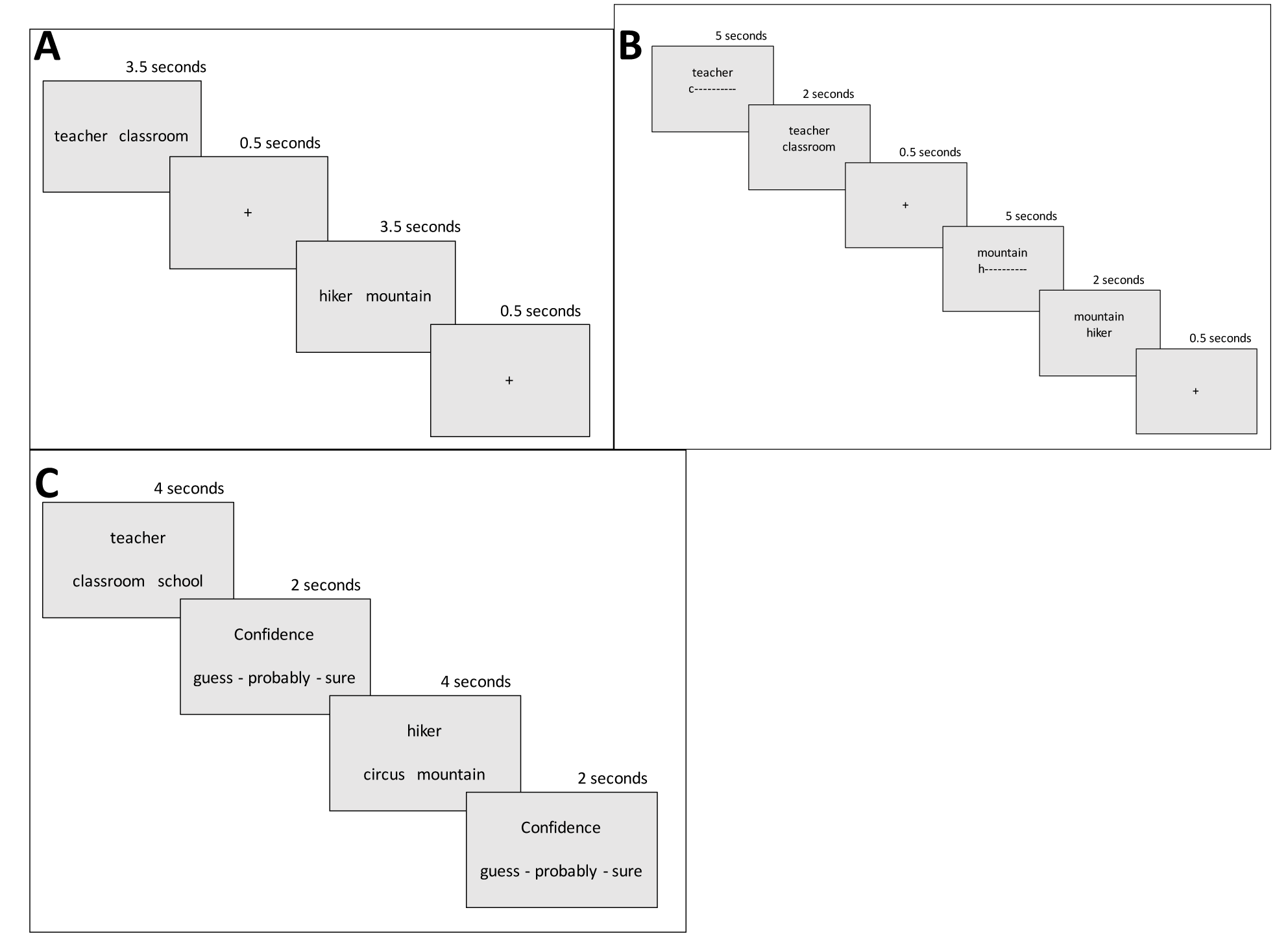
Representations of trial procedure for study phase **(A)**, cue-recall learning task **(B)**, and test phase **(C).**

#### Cue-recall learning task

Following the study phase, in order to saturate learning for both schema-consistency conditions, participants performed multiple drop-out cycles of cued-recall of all studied pairs. In each cycle, participants were presented with all cue words (one of the words they had previously studied) in a random order, and asked to type in the word that had been paired with each cue. Participants were presented with the first letter of the correctly matching word, and had five seconds to type the remaining letters. After typing in a word, participants were shown the correct answer paired with the cue word (regardless of accuracy), to enable restudying. In the case when participants typed an incorrect word as the match for the cue word, the cue word was added to the end of the list and was tested again following the presentation of all other cue words. This continued until all pairs were correctly recalled, thus concluding one learning cycle. During the following cycles, the cue words alternated in presentation order: first round presented with the person and required to type in the matching location, second round presented with the location and required to type in the matching person. The task completed once each pair had been recalled correctly three consecutive times without drop-out.

#### Distractor task

Once the cue-recall task was completed participants played a game of Tetris (http://www.freetetris.org/game.php) for 15 minutes. This distractor task allowed us to eliminate the recently learned information from working memory, and to prevent rehearsal of the word pairs.

#### Forced-choice test phase

During the test phase, on each trial participants saw three words, a cue word on top and two choices on the bottom. Participants completed a forced-choice task by selecting which of the two words on the bottom had been associated with the cue word *within the experiment ‐* either because they were studied together or because they were indirectly connected by studied pairs. To prevent participants from responding solely based on schema consistency in the schema consistent condition, the foil was selected to be as strongly semantically related to the cue (*M_LSA_* = .28, *SE* = .02), as was the correct answer (*M_LSA_* = .27, *SE* = .02). E.g., if the cue was “teacher” and the correct answer was “classroom” the distractor was “school”. Words from the schema inconsistent condition served as foils for the schema consistent condition and vice versa. Therefore, all words in the forced-choice recognition task were encountered in the study phase, and participants could not respond based on familiarity alone.

Participants were instructed to answer as quickly and as accurately as possible, as both factors would increase the amount of points they earned for the task. The points (later displayed to the participants) were helpful for keeping participants motivated, without causing additional learning during the test phase (because the point-feedback was not provided on the basis of individual trials). There were two ways in which correct words could be associated with the cue word: i) direct associations occurred when the two words (i.e. A-B) had been seen previously during the study phase and recognition task; ii) indirect associations occurred when words had been learned, but never directly paired together (i.e. A-C, since previously learned A-B and B-C). Participants were not required to make the distinction between direct and indirect associations, but instead simply selected which word was in some way associated with the cue word. After selecting a word, participants were asked to indicate their confidence in their answer on a 3-point scale (guess, probably, sure). After 10 trials, participants were shown a screen with the number of points they had accrued up to that point and could rest if needed, before beginning a new set of trials.

There were 100 test trials: 60 containing studied pairs and 40 containing linked pairs. In order to gather more observations, the testing was repeated 4 times, where each cycle contained the 100 trials we described in a novel random order each time. Each pair was tested in both directions twice ‐the person was the cue for two of the trials, while location was the cue for the remaining two. After completing the four test cycles, participants completed a brief questionnaire and were compensated for their time.

## RESULTS

We analyzed the accuracies, confidence ratings, and RTs via logistic and linear mixed-effects regression models (Baayen, Davidson, & Bates, 2008). We excluded incorrect responses from analyses of confidence ratings and RTs (6-30%, depending on the condition). Random effects were determined through restricted likelihood ratio tests and all final models included varying intercepts for subjects and individual word pairs (i.e., different subjects and items differ in their overall accuracy and RT estimates), as well as varying slopes by subject for the effect of schema consistency (i.e., the models account for how much differences in schema consistency varies across subjects). We inferred the significance of each effect based on likelihood ratio tests and AIC comparisons of the regression models that contained the effect in question with identical models that lacked this contrast.

### The effect of schema on learning associations

During the study phase, initial learning differed between schema-consistent pairs and schema-inconsistent pairs. Schema-consistent word pairs were correctly recalled more often on their first presentation in each cued-recall cycle (Figure 2a; *ΔAIC* = −34, *χ^2^* (1) = 35.82, *p* < .001). Schema-consistent pairs were also recalled faster (Figure 2b; *ΔAIC* = −26, *χ^2^* (1) = 27.62, *p* < .001) and with higher accuracy throughout the study phase (Figure 2c; *ΔAIC* = −37, *χ^2^* (1) = 38.97, *p* < .001), though memory for the pairs was saturated in both conditions by the end of learning. This is evident by the subsequent forced-choice recognition performance for studied word pairs (Figure 3), which were recognized equally accurately (*ΔAIC* = −0.7, *χ^2^* (1) = 2.69, *p* = .101) and with similar confidence (*ΔAIC* = 0.8, *χ^2^* (1) = 0.229, *p* = .632) regardless of schema consistency. Schema-consistent pairs were recognized slightly faster than schema-inconsistent pairs, but the effect did not reach significance (*ΔAIC* = −1, *χ^2^* (1) = 3.41, *p* = .065). In summary, while it took longer to learn schema-inconsistent pairs to criterion, post-learning recognition accuracy, speed and confidence did not differ as a function of schema consistency.

**Figure 2.**
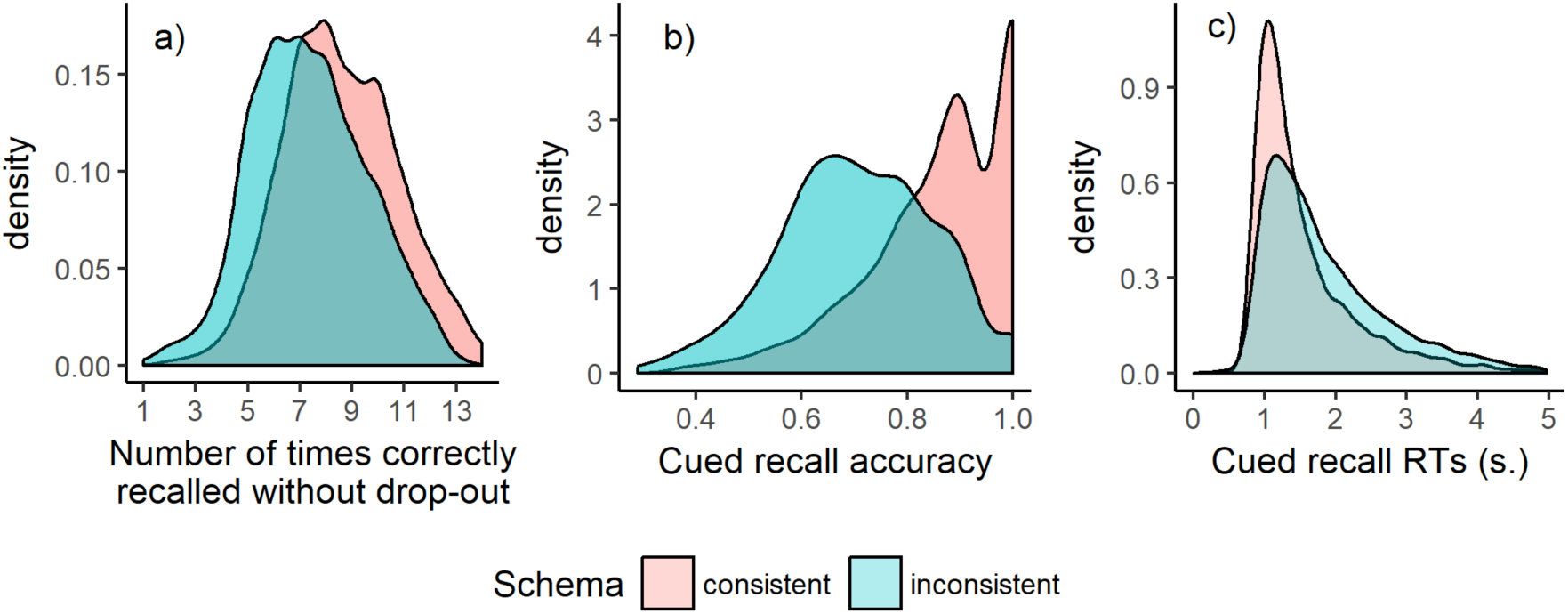
Cued-recall performance during learning: a) distribution of the number of cued-recall cycles on which each pair was recalled correctly on the first presentation (i.e. without further drop-out); b) distribution of cued-recall accuracy for each pair averaged over learning cycles; c) distribution of RTs for correct cued-recall of each pair.

**Figure 3.**
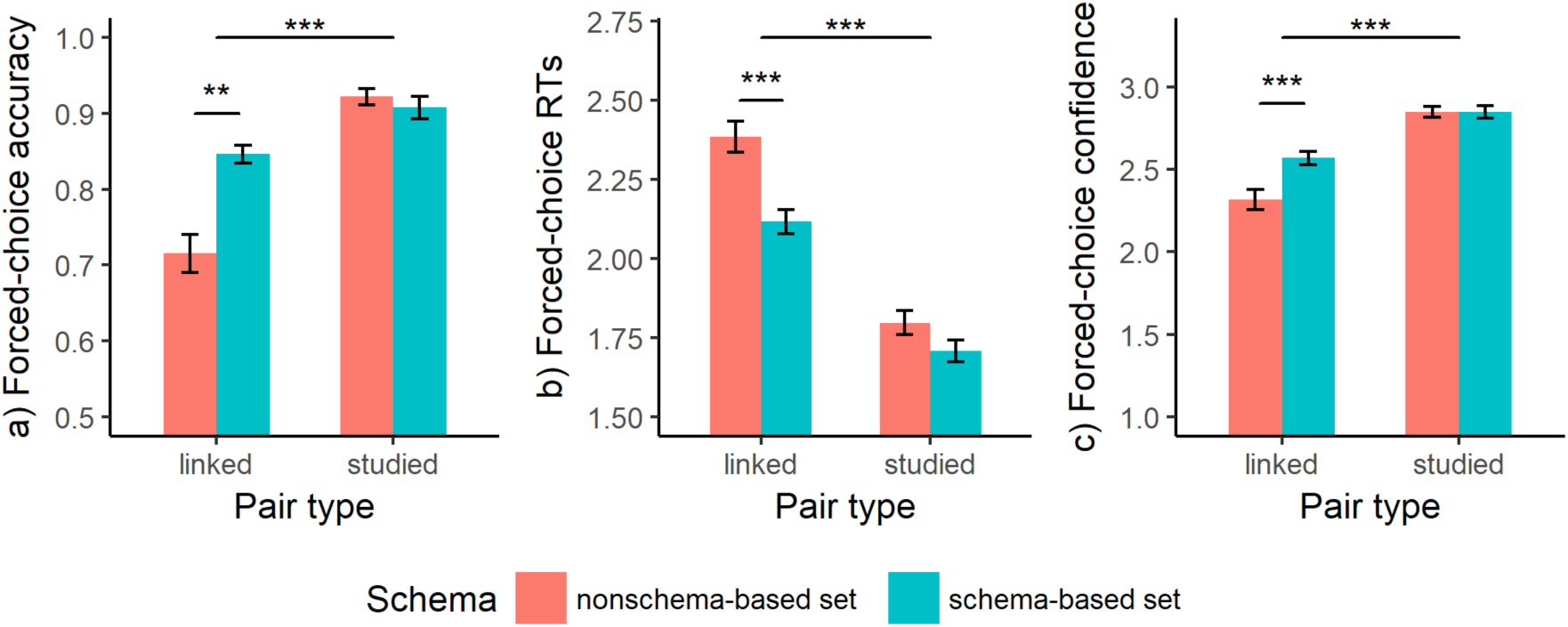
Forced choice a) accuracy, b) RTs (in s.) and c) confidence for studied and linked pairs depending on whether they were part of a schema. ^**^ *p* < .01, ^**^ *p* < .001

### The effect of schema on memory integration

Forced-choice performance in the recognition test of “linked” pairs (A-C or A-D for A-B, B-C, C-D) revealed that there was greater memory integration for pairs that were in schema-consistent sets. Participants were more accurate (Figure 3a, *ΔAIC* = −8, *χ^2^* (1) = 10.6, *p* = < .01), faster (Figure 3b, *ΔAIC* = −14, *χ^2^* (1) = 15.68, *p* < .001) and more confident (Figure 3c, *ΔAIC* = −59, *χ^2^* (1) = 65.49, *p* < .001) in judging that pairs had been linked, when they were part of a schema. This is despite accuracy and confidence of studied pairs being saturated by the end of the study phase. In summary, all three measures of forced-choice performance (accuracy, RTs and confidence) indicate that schema-consistent linked pairs were better integrated during study.

Could the differences in forced-choice performance for linked pairs be explained by the speed of learning (i.e., the number of study trials to reach criterion)? Memory integration involves reactivating traces for related information (Karlsson and Frank, 2009; Shohamy and Wagner, 2008) and since schema-consistent pairs were learned faster and earlier during the study, they might be easier to reactivate and integrate. To test this explanation, we included the average accuracy and RTs for each same-set word pair for each participant, as predictors and by-subject random slopes in the mixed-effects regression model. Memory integration was better when the component pairs were learned more quickly during study – average cued recall accuracy for studied pairs predicted subsequent forced-choice accuracy (*ΔAIC* = −10, *χ^2^* (1) =12.387, *p* < .001) and confidence ratings (*ΔAIC* = −23, *χ^2^* (1) =24.65, *p* < .001) for linked pairs in the set, and the cued recall RTs of studied pairs predicted forced-choice RTs for linked pairs in the set (*ΔAIC* = −4, *χ^2^* (1) =6.21, *p* = .013). Importantly, the differences between schema-consistent and schema-inconsistent pairs that we outlined above remained significant even after accounting for the learning rate of each pair (all *p* < .01), suggesting that learning rate was not driving this effect.

### The effect of schema on integrative encoding

In the last section, we presented evidence that schema-consistent pairs experience more memory integration. This section further looks into the extent of fast memory integration (i.e. integrative encoding), which is the focus of the current study. All three measures of forced-choice performance indicate there was more integrative encoding for schema-consistent pairs. The extent of integrative encoding during memory integration is reflected in the independence of accuracy, RTs and confidence from the number of links among the linked pairs. If participants depended more on logical inference to judge the pairs, then accuracy, RTs, and confidence should have been worse when they had to cognitively traverse more links to connect the words. Table 2 shows that the number of links impacted all measures twice as much for schema-inconsistent, than schema-consistent, pairs. The mixed effects regression models confirmed a significant main effect of number of links on accuracy (*ΔAIC* = −26, *χ^2^* (1) = 28.56, *p* < .001), RTs (*ΔAIC* = −19, *χ^2^* (1) =21.15, *p* < .001) and confidence (*ΔAIC* = −22, *χ^2^* (1) = 24.30, *p* < .001). The effect of number of links interacted significantly with schema consistency for RTs (*ΔAIC* = − 3, *χ^2^* (1) = 5.04, *p* = .025) and confidence (*ΔAIC* = −7, *χ^2^* (1) = 8.94, *p* = .003), but not accuracy (*ΔAIC* = 0.7, *χ^2^* (1) = 1.26, *p* = .261). These results are consistent with successful memory integration requiring integrative encoding in the schema-consistent condition, while decisions in the schema-inconsistent condition depend on logical inference.

**Table 2.** Forced-choice performance.

| Condition | Accuracy | RTs (ms.) | Confidence |
| --- | --- | --- | --- |
| Schema consistent |  |  |  |
| Links: 1 | 0.88 | 2064 | 2.79 |
| Links: 2 | 0.81 | 2163 | 2.61 |
| **Links: difference** | **0.07** | **-99** | **0.18** |
| Schema inconsistent |  |  |  |
| Links: 1 | 0.79 | 2268 | 2.67 |
| Links: 2 | 0.65 | 2485 | 2.30 |
| **Links: difference** | **0.14** | **-217** | **0.37** |

## GENERAL DISCUSSION

We have conducted a study in which participants learned pairs of words with overlapping content (A-B, B-C, C-D) during a study phase. Shortly after a distraction task, participants judged whether two elements were indirectly linked during the study phase (A-C or A-D). We varied the relation of word pairs to be either schema-consistent or schema-inconsistent. The results of this study confirmed our hypothesis that schema not only affects the initial learning of word associations, but also affects performance in later memory integration (even after accounting for differences in initial learning rate). In addition, the data suggest that schema consistency facilitates integrative encoding through a weaker dependency of inference RTs, accuracy and confidence on the number of links that connected the two words during study.

### Extending the effect of schema to memory integration

It is known that schema affects the initial acquisition, and consolidation, of memories (Alba and Hasher, 1983). New information can undergo system-level consolidation (with hippocampal independence) very rapidly when facilitated by a schema (Tse et al., 2007). In fact, even among patients with medial temporal lobe (MTL) damage, intact prior knowledge structures can support learning new episodic information that is consistent with schemas (Kan, Alexander, & Verfaellie, 2009). In contrast, damage to the mPFC is associated with reduced ability to integrate incoming information (Schnider, 2003). Recent neural imaging studies in a healthy population further verified that schema-consistent knowledge is mediated by mPFC, and is integrated with neocortex rapidly, while schema-inconsistent knowledge is mediated by the MTL (van Kesteren et al., 2012; van Kesteren et al., 2013). These differences in initial acquisition and consolidation of new memories motivated us testing for memory integration in our current study.

In particular, we observed not only enhanced learning of word pairs facilitated by schema, but also enhanced memory integration, in the form of improved recognition of schema-consistent linked pairs. This enhanced memory integration occurred despite both schema-consistent and inconsistent studied pairs being learned to the same criterion, and recognized equally well during the test phase. In addition, the facilitation in memory integration is almost immediate during the encoding stage, rather than occurring (through logical inference) during the retrieval of initially learnt word pairs. When combined, these results suggest that schema plays a key role in fast integration of new information with existing memories.

### Potential mechanism of fast memory integration facilitated by schema

Given the differential involvement of the hippocampus or neocortex based on the use of schema (van Kesteren, Ruiter, Fernandez, & Henson, 2012; van Kesteren et al., 2013), we propose that the facilitation effect that we observed is a result of different mechanisms of memory integration in the hippocampus and neocortex.

In the hippocampus, fast memory integration is facilitated by dynamic shifts between encoding and retrieval states. Encoding of a new but overlapping event can reactivate a previous event that has mismatching details (Karlsson and Frank, 2009; Shohamy and Wagner, 2008). In the neocortex, gradual generalization across overlapping information is a known possibility (McClelland et al., 2010; Rogers and McClelland, 2004; McClelland and Rogers, 2004), but rapid integration requires fast initial encoding of information. Fast encoding in the neocortex is possible when the new information is consistent with schema (Tse et al., 2007, 2011), which has been associated with the mPFC inhibiting hippocampal binding (van Kesteren, Ruiter, Fernandez, & Henson, 2012; van Kesteren et al., 2013).

One question for future research is why fast memory integration did not occur in the schema-inconsistent condition. It is possible that the generalization mechanism is more efficient in the neocortex than in the hippocampus, and that fast memory integration in the hippocampus can still occur but requires more training and exposure to the material. Alternatively, facilitating the integration of schema-inconsistent information might depend on the type of novelty that the hippocampus detects: whether new material is unrelated to, or incongruent with, the existing schema (van Kesteren et al., 2012; Kumaran and Maguire, 2009). It might be possible to observe fast memory integration with a set of stimuli that defines schema-inconsistency differently.

### Further implications and future directions

The current study suggests an alternative mechanism that supports integrative encoding. In particular, when there is rapid system-level consolidation facilitated by schema, memories can also undergo fast memory integration. Given the short interval between our study and test phases, it is likely that the learnt material had not become completely independent of the hippocampus, even under the effect of schema. The resulting integrative encoding is therefore likely supported by both the hippocampus and neocortex, or from an interaction between the two. Future neural imaging studies with high spatial resolution in different cortical and sub-cortical areas would be beneficial to make this distinction. Further neural imaging studies might also use neural activity during the encoding stage to predict later memory integration performance.

## ACKNOWLEDGEMENTS

We thank John Paulus, Kimberly Hoover and Chao Wu for their assistance with the study.

## Appendix A

Full set of experimental stimuli.

| Condition | Stimuli type |  |  |  |
| --- | --- | --- | --- | --- |
|  | Person 1 | Location 1 | Person 2 | Location 2 |
| Schema consistent | Teacher | Classroom | student | Dormitory |
|  | Doctor | Hospital | Patient | Reception |
|  | Judge | Courtroom | Lawyer | Firm |
|  | Astronaut | Spacecraft | Pilot | Cabin |
|  | Fireman | Station | policeman | precinct |
|  | Hunter | Forest | Ranger | Park |
|  | Chef | Kitchen | Waiter | Parlor |
|  | Player | Arena | Announcer | Newsroom |
|  | Bellhop | Hotel | Guest | Mansion |
|  | Driver | Road | Racer | Stadium |
| Schema inconsistent | Baker | Mountain | Principal | Circus |
|  | Scientist | Street | Nurse | Tavern |
|  | Cyclist | Inn | Prosecutor | Library |
|  | Maid | Track | Explorer | Casino |
|  | Surgeon | School | Officer | Lodge |
|  | Guard | Restaurant | Chief | Ballroom |
|  | Professor | Rocket | Hostess | City |
|  | Gladiator | Chamber | Coach | Ward |
|  | Lumberjack | Highway | Clerk | Airport |
|  | Bailiff | Pharmacy | Motorist | Office |

